# The genetic signature of memory encoding along the human hippocampal axis

**DOI:** 10.64898/2026.06.22.733862

**Authors:** Mantre Dehnad, Tung To, Haley Moore, Anne Freelin, Ashwinikumar Kulkarni, Stephan Loer, Bradley Lega, Genevieve Konopka

**Affiliations:** Department of Neuroscience, University of Texas Southwestern Medical Center; Dallas, TX, 75390, USA; O’Donnell Brain Institute, University of Texas Southwestern Medical Center; Dallas, TX, 75390, USA; Department of Anesthesiology, Amsterdam University Medical Center; Amsterdam, 1081HV, The Netherlands; Department of Neurosurgery, University of Texas Southwestern Medical Center; Dallas, TX, 75390, USA; Department of Neurobiology, University of California Los Angeles; Los Angeles, CA, 90095, USA

## Abstract

Episodic memory formation engages hippocampal oscillations that vary along the anterior-posterior axis, but the molecular programs supporting this physiological specialization remain unclear. Here, we leveraged a rare neurosurgical dataset in which patients performed verbal episodic memory tasks during intrahippocampal intracranial EEG recordings prior to en bloc hippocampal resection, enabling integration of encoding-related oscillatory signatures with matched cell-type-resolved transcriptomics from the same individuals. Subsequent memory effects (SMEs) spanned delta/theta, gamma, and hippocampal ripple activity across anterior and posterior hippocampus. Single-nucleus RNA sequencing from anatomically matched anterior and posterior tissue revealed longitudinal transcriptional gradients, most prominent in excitatory neurons. Spatial transcriptomic maps validated axis-enriched transcripts and their localization. Linking subject-specific SMEs to gene expression identified distinct molecular programs: anterior low frequency SMEs associated with synaptic and chromatin-regulatory pathways, and posterior high-frequency SMEs associated with metabolic and protein synthesis processes. Gene regulatory network inference further revealed axis-specific hub architectures. Together, these results define a cell-type-specific genetic architecture linking longitudinal molecular specialization to the human hippocampal encoding dynamics.

## Introduction

Episodic memory enables humans to remember events embedded in their spatio-temporal contexts and to recall these experiences to guide future behavior and decision-making^1–3^. Within the medial temporal lobe (MTL), the hippocampus plays a critical role in the formation and retrieval of such episodic memories by indexing localized cortical neuron populations to bind the elements of an experience into a coherent memory^2, 4, 5^. Intracranial recordings in human neurosurgical patients have revealed key oscillatory activity relevant to memory encoding, including delta (∼2-5Hz, slow theta), theta (∼5-9Hz, fast theta), low- (30-70Hz) and high-gamma (70-120Hz) rhythms^6–9^, as well as brief high-frequency ripple events^10–12^. In the human hippocampus, mnemonically-relevant activity occurs across the 2-10 Hz range, with dissociable properties for both fast (5-9 Hz) and slow (2-5 Hz) activity. Changes in oscillatory power, and connectivity, especially in the theta frequency range, reliably predict successful item encoding, while ripple activity has been linked to consolidation and replay processes. These physiological signatures arise within a hippocampal circuit that is itself not uniform along its longitudinal axis.

Previous electrophysiological, imaging, structural, and genetic studies have demonstrated distinct functional specializations along the longitudinal (anterior-posterior, A-P) axis of the human hippocampus^13^. The anterior hippocampus (aHC) is more tightly coupled to affective and motivational circuits and supports gist-like integration, whereas posterior (pHC) areas interact more with the posterior cortex and support detailed spatial and contextual information^14–19^. Furthermore, human iEEG studies have established axis-specific gradients during memory tasks, especially in delta/slow theta and (5-9 Hz) fast theta rhythms, and hippocampal ripple events that emerge preferentially during successful encoding and are distinct from sleep associated sharp-wave ripples^20–27^. Together, these observations indicate that the physiology supporting episodic encoding varies systemically along the hippocampal axis. The transcriptional programs that underly these memory-relevant spatial differences remain largely unknown, despite their importance toward understanding the molecular basis of memory formation. Such data can reveal mechanisms by which local circuit are tuned for distinct mnemonic functions and may provide targets for interventions aimed at enhancing memory or ameliorating memory impairments.

snRNA-seq provides the molecular resolution to dissect these gene expression programs, but obtaining suitable human hippocampal tissue specimens following behavior is a major challenge. We previously compared anterior and posterior hippocampal tissue obtained from en bloc surgical resections to delineate cell-type specific transcriptional differences that suggested intrinsic molecular specialization^28^. Complementing these findings in the hippocampus, our prior studies in the neocortex demonstrated that band-limited oscillations linked to SMEs map onto gene sets enriched for synaptic, chromatin, and metabolic pathways^29, 30^. This prior study demonstrated that single-cell molecular profiles of human brain tissue could be integrated with oscillatory dynamics to resolve memory-specific effects.

Here, we link data describing spatially precise hippocampal oscillatory dynamics of memory encoding with single-cell gene expression profiles to investigate the transcriptional programs that regulate encoding-related activity. Subjects performed episodic memory tasks during iEEG, allowing us to generate oscillatory fingerprints of successful encoding, including theta oscillations and ripple events. We then combine these data with cell-type resolved snRNA-seq from anatomically matched anterior and posterior hippocampal tissue from *the same individuals*. This allows us to directly test whether longitudinally-specific oscillatory signatures, spanning delta, theta, gamma and hippocampal ripple activity, align with gene expression programs within defined excitatory and inhibitory neuronal populations, and to extend this to non-neuronal cell types. Using this novel platform, we demonstrate that the oscillatory dynamics and gene expression programs vary together across the hippocampal longitudinal axis, paralleling the region’s physiology. By integrating memory-related physiology, transcriptomics, spatial validation, and gene-regulatory network inference, we define a genetic architecture that links longitudinal molecular specialization across cell classes to the neurophysiology of human hippocampal encoding.

## Results

### Linking memory-related hippocampal physiology to molecular specialization along the longitudinal axis

To characterize the physiological signatures of successful episodic encoding along the hippocampal axis, we analyzed iEEG from twelve patients undergoing presurgical evaluation for intractable temporal lobe epilepsy; one patient did not meet iEEG quality criteria and was excluded from electrophysiological analyses, leaving eleven patients for SME quantification (Supplementary Table S1). Participants completed Associative Recognition and/or Free Recall episodic memory tasks during continuous monitoring (Fig. 1A, see Methods). Depth electrodes sampled both anterior and posterior hippocampus simultaneously (aHC: 34 contacts, pHC: 26 contacts), with expert neuroradiologically-confirmed anatomical localization.

**Figure 1.**
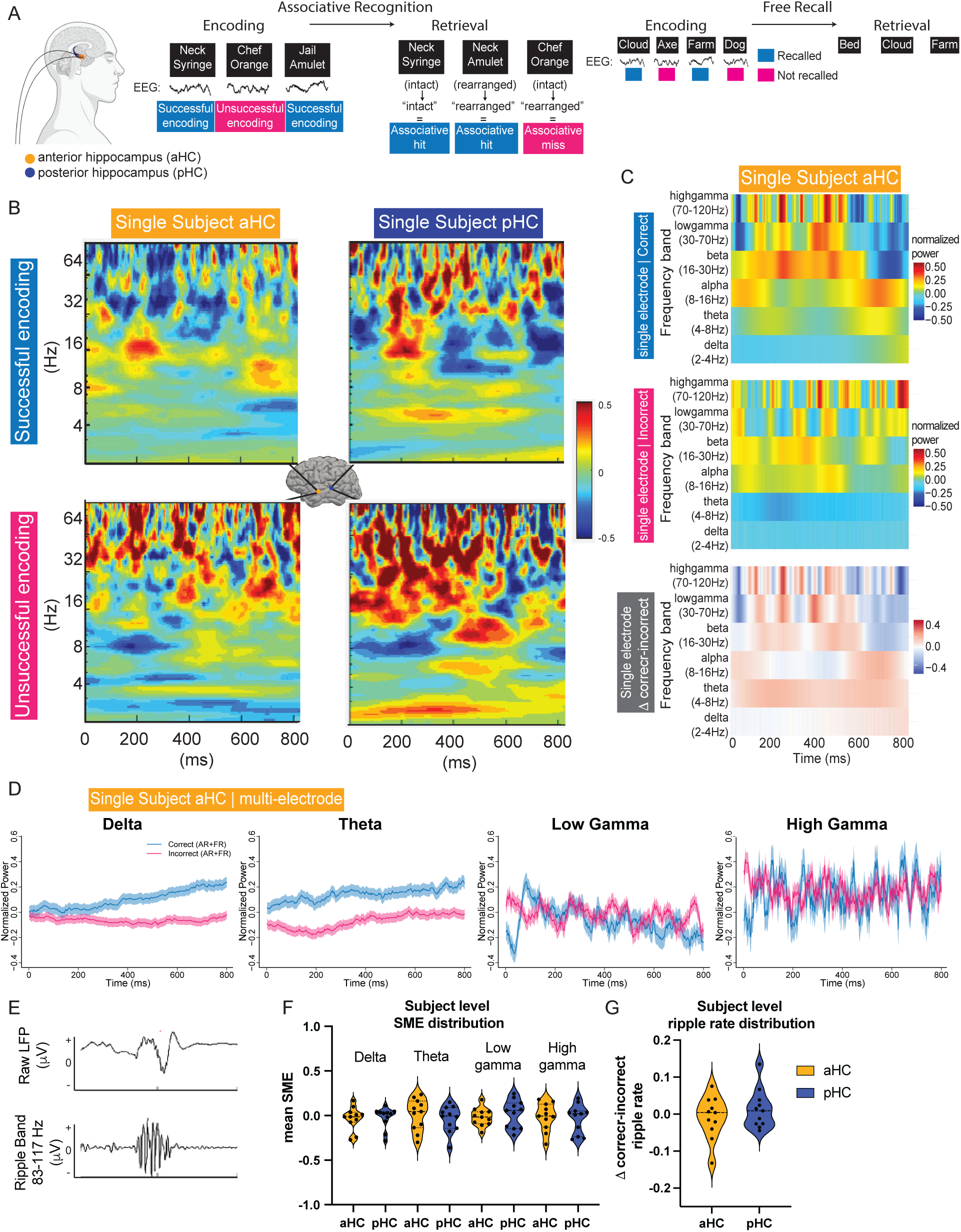
Oscillatory signatures of episodic memory encoding across the hippocampal axis. **A:** Left: Graphical representation of relative implantation region of iEEG in aHC and pHC during episodic memory testing. Middle: Associative Recognition (AR) paradigm; correct encoding yields “associative hit,” incorrect yields “associative miss.” Right: Free Recall (FR) paradigm; correct encoding yields “recalled,” incorrect yields “not recalled.” **B:** Example of a singular aHC and pHC depth-electrode location and corresponding time-frequency plots for correct and incorrect encoding. Y-axis shows frequency (Hz), X-axis time (ms). Power is baseline-normalized (−0.5-0 s) and z-scored. **C:** Compression of z-scored power in defined oscillatory bands in delta (2–4 Hz), theta (4–8 Hz), alpha (8–16 Hz), beta (16–30 Hz), low-γ (30–70 Hz), high-γ (70–120 Hz) frequencies. During the first 800ms post-stimulus, for correct (top) and incorrect (middle) encoding, and difference of normalized power between correct and incorrect encoding trials (bottom) **D:** SMEs were computed per subject (*s*), region (*r*), and per oscillatory band (*f*), as the power difference between correct and incorrect trials, averaged across all electrodes in the aHC or pHC, in memory relevant oscillatory bands. **E:** Schematic of ripple band detection, top: raw LFP trace during onset of stimulus, bottom: LFP filtered for high frequency ripple (adapted from Seger et al 2024) **F:** Distribution of subject level (individual dots) SMEs across memory relevant oscillations across anterior and posterior hippocampus, **G:** Difference between subject level mean ripple rate during correct and incorrect trials. (humanoid graphical in panel A created in https://BioRender.com)

Spectral power was extracted during the first half of the encoding epoch following stimulus onset in delta (2-4Hz), theta (4-8Hz), alpha (8-16Hz), beta (16-30Hz), low-gamma (30-70Hz), high-gamma (70-120Hz) bands (Fig. 1B, C), a time frame previously shown to exhibit memory-related oscillatory signatures across paradigms^22^. We also extracted discrete ripple events during recall using consensus criteria for human data (Fig. 1E). Example electrodes illustrate the differences across multiple frequency bands.

SMEs were quantified as the difference in oscillatory power between remembered (“recalled/correct”) and forgotten (“nonrecalled/incorrect”) trials, computed per subject, region, and oscillatory band (Fig. 1D, F), and as the difference between mean ripple rate between remembered and forgotten trials (Fig. 1E, G). Across subjects, successful encoding was associated with band-limited SMEs spanning delta, theta and gamma frequencies, as well as changes in ripple rate, quantified at the subject level for each hippocampal pole (Fig. 1F-G). These subject-level physiological features provided the basis for subsequent alignment with subject-level molecular profiles obtained from matched anterior and posterior hippocampal tissue (Fig. 2A-D, Supplementary Fig. S1, Supplementary Table S2).

**Figure 2.**
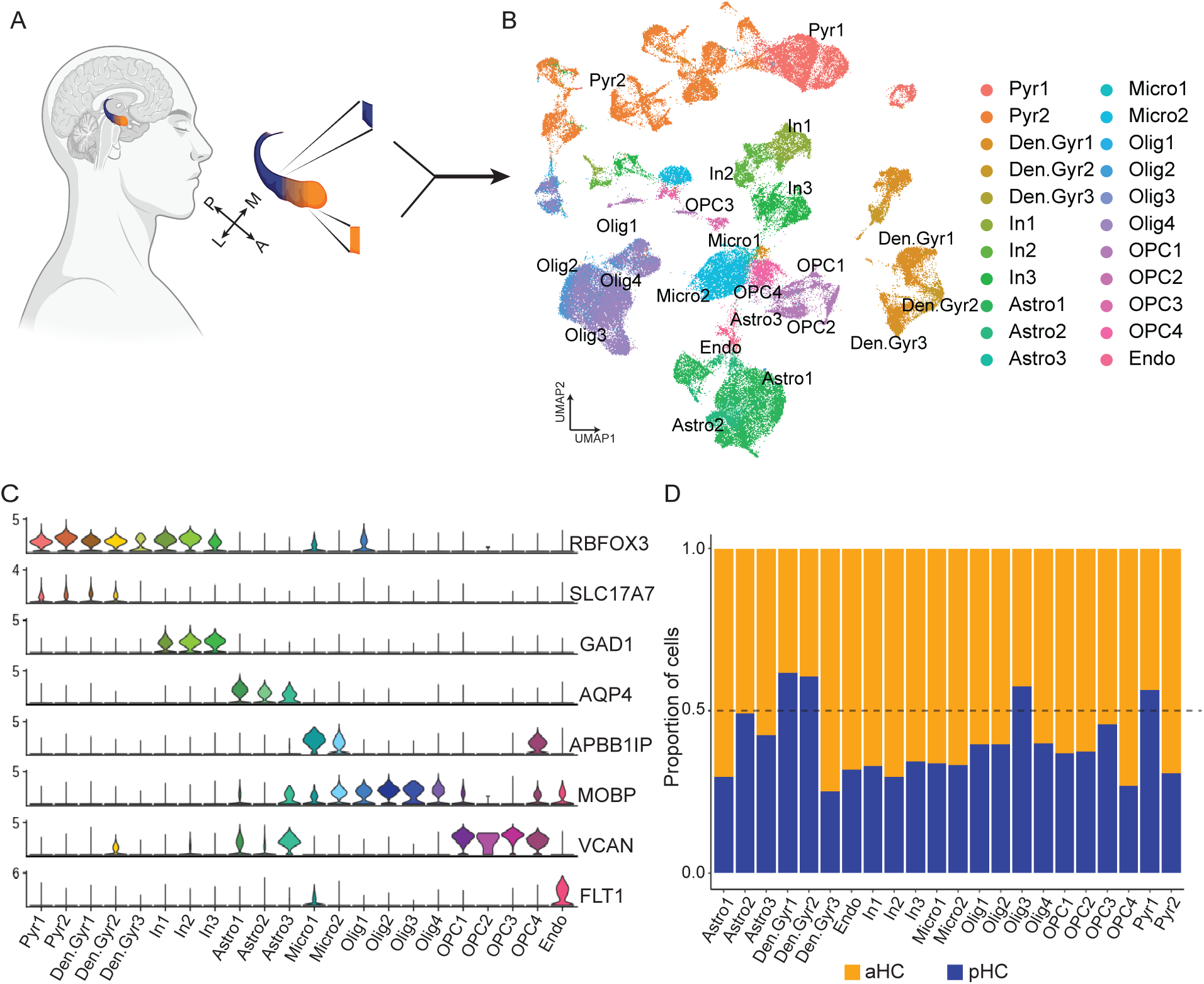
Single-nucleus RNA sequencing of anterior and posterior human hippocampus. **A:** En bloc resection of the ipsilateral hippocampus (∼3.5 cm) permitted separation of anatomically defined anterior (aHC; proximal ∼0.5 cm) and posterior (pHC; distal ∼0.5 cm) tissue. Twelve patients contributed paired aHC/pHC samples (24 total). **B-C:** Nuclei were isolated and profiled using the ScaleBio snRNA-seq platform. After QC filtering, UMAP embedding revealed all major neuronal and glial populations, annotated using canonical markers and verified using our published hippocampal atlas (Ayhan et al., 2021). **D:** aHC and pHC proportional contribution to all major cell classes (Graphical in panel A created in https://BioRender.com)

Matched anterior and posterior hippocampal tissue from the same individuals enabled testing whether these physiological encoding signatures aligned with underlying molecular specialization. Hippocampi were resected using an en bloc technique, permitting spatially precise sampling along the longitudinal axis. Only hippocampi of minimum 3.5 cm in length (after removal) without evidence of radiographic abnormality (including mesial temporal sclerosis, MTS, tumors, or vascular lesions) were included for transcriptomic analysis. Anterior and posterior samples were taken from the proximal and distal ends of each specimen corresponding to locations approximately 1.5 cm in front or behind the uncal notch, respectively. These locations were selected to match implantation targets during presurgical intracranial mapping.

Resected hippocampal tissue from twelve patients (Supplementary Table S1) was processed for snRNA-seq (see Methods). Integration with our previously published human hippocampal reference dataset^28^ and validation using canonical marker genes resolved all major neural and glial classes (Fig. 2A-D; see Methods). All major cell-types were consistently observed in both anterior and posterior hippocampal samples across donors (Fig. 2D), enabling region-specific comparisons while accounting for inter-individual variability.

To assess whether transcriptional programs differ along the hippocampal axis within this cohort, independent of oscillatory activity, we first focused on the neurons in these regions (*RBFOX3* positive cells). We identified all subtypes of excitatory and inhibitory neurons using *SLC17A7* and *GAD1* as broad marker genes, respectively (Fig. 3A-B, Supplementary Fig. S2). Combining nuclei across these major annotations of either excitatory or inhibitory neurons, we carried out pseudobulk differential expression analyses comparing anterior and posterior hippocampal nuclei and found pronounced anterior-posterior transcriptional differences (Fig. 3 C, D).

**Figure 3.**
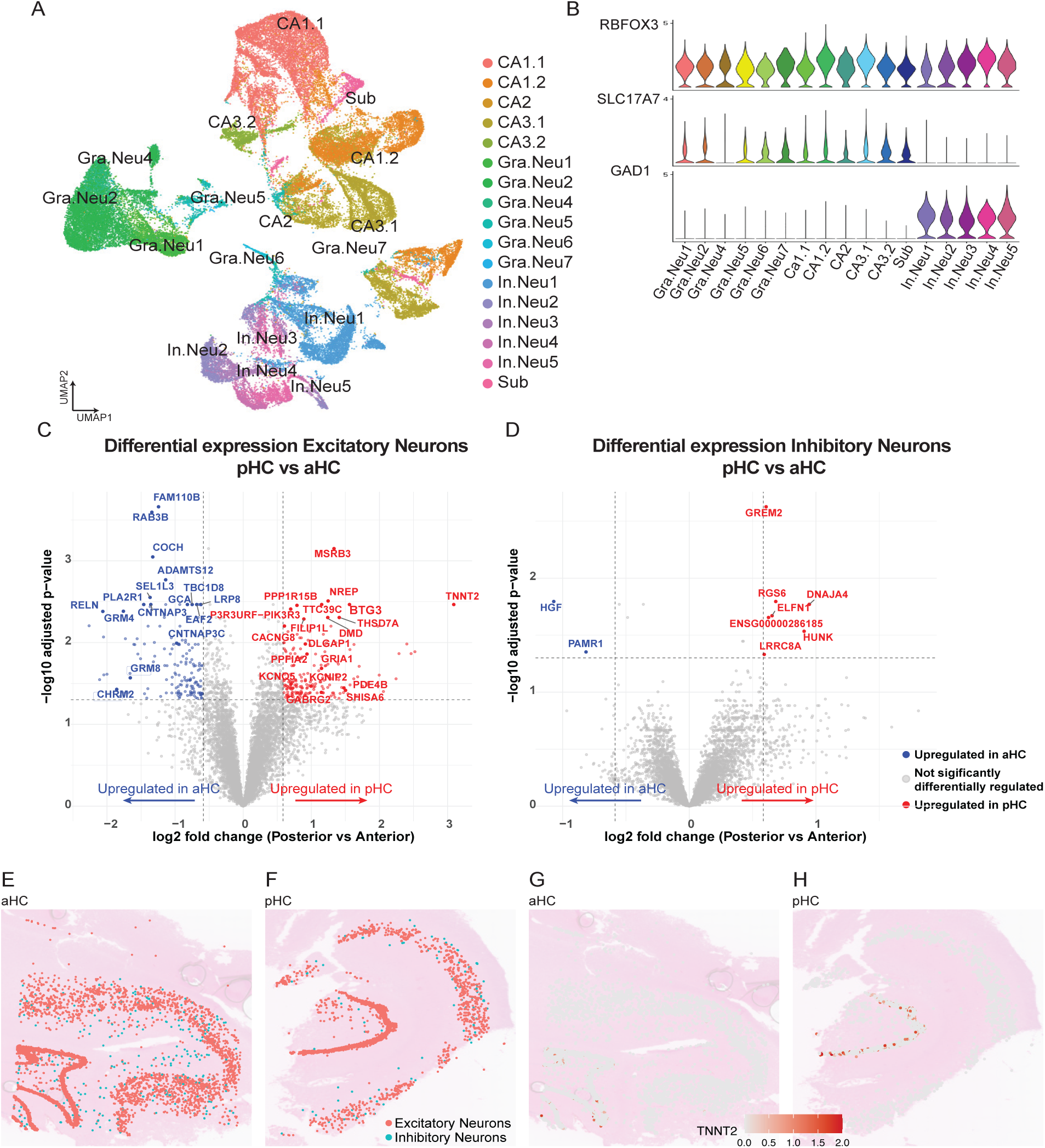
Cell type–specific transcriptional gradients validated by spatial transcriptomics. **A-B:** Subclustering of excitatory and inhibitory neuronal subclasses after integration with our reference dataset (Ayhan et al. 2021) highlighting expected laminar and subtype diversity. **C:** Volcano plot for posterior > anterior differential expression within excitatory neurons (DESeq2 pseudobulk; covariates: *SeqBatch*, sex, age, epilepsy duration, hemisphere. Significance log2FC ± 0.585, -log10(FDR <0.05). Top 10 anterior- and posterior-enriched genes are labeled. Posterior-enriched genes include *TNNT2* (log2FC = 3.09, FDR-*q* = 0.0034), *GRIA1* (1.15, *q* = 0.0207), *GABRG2* (1.15, *q* = 0.0406), *PDE4B* (1.48, *q* = 0.036), *DLGAP1* (0.91, *q* = 0.010), *PPFIA2* (0.94, *q* = 0.0138), *SHISA6* (1.50, *q* = 0.038), and *CACNG8* (0.60, *q* = 0.0063); anterior-enriched examples include *RELN* (−2.06, *q* = 0.0042), *GRM4* (−1.76, *q* = 0.0044), *GRM8* (−1.66, *q* = 0.027), and *CHRM2* (−1.86, *q* = 0.037). **D:** Volcano plot for inhibitory neurons highlighting posterior enrichment of *GREM2* (0.61, *q* = 0.0024), *ELFN1* (0.65, *q* = 0.021), *RGS6* (0.68, *q* = 0.016), *DNAJA4* (0.95, *q* = 0.017), and anterior enrichment of *HGF* (−1.07, *q* = 0.016). **E-F:** Visium HD spatial maps from aHC (E) and pHC (F) slices showing major neuronal cell types. **G-H:** Spatial feature plots for *TNNT2* demonstrate stronger expression in excitatory cells in the pHC (H) relative to aHC (G), with signal concentrated in the dentate gyrus.

Excitatory neurons, including pyramidal and dentate gyrus populations, showed 358 differentially expressed genes (DEGs) between anterior and posterior hippocampus (Supplementary Tables S3 and S4). Posterior excitatory neurons were enriched for genes involved in synaptic transmission, dendritic signaling, and postsynaptic organization, including *GRIA1*, *GABRG2*, *DLGAP1*, *PPFIA2*, *SHISA6*, *PDE4B*, *CACNG8*, *KCNQ5*, and *KCNIP2* (Fig. 3C). In contrast, anterior excitatory neurons were enriched for *RELN*, *GRM4*, *GRM8*, *CHRM2*, *CNTNAP3*, and *CNTNAP3C*, indicating increased representation of metabotropic, modulatory, and adhesion-related signaling pathways (Fig. 3C).

Inhibitory neurons showed more modest anterior–posterior transcriptional differentiation, with 9 DEGs identified between the two regions (Supplementary Tables S3 and S4). Posterior inhibitory populations were enriched for genes involved in synaptic coupling, presynaptic release dynamics, and G-protein-dependent ion channel regulation, including *GREM2*, *ELFN1*, *RGS6*, and *HUNK* (Fig. 3D). Anterior inhibitory neurons preferentially expressed *HGF*, a regulator of interneuron maturation and excitability. Together, these findings indicate that anterior–posterior transcriptional specialization is present in both excitatory and inhibitory neuronal populations.

To validate these anterior-posterior transcriptional gradients in situ, we generated high-resolution Visium HD spatial transcriptomic maps from independent anterior and posterior hippocampal resections (Fig. 3E-F; see Methods). Spatial clustering recapitulated the major neuronal and glial classes identified by snRNA-seq (Supplementary Fig. S3), and region-enriched transcripts showed concordant spatial localization. For example, *TNNT2*, a posterior-enriched excitatory marker (Fig. 3C), exhibited higher expression in posterior hippocampus with prominent localization to the dentate gyrus (Fig. 3G-H), consistent with snRNA-seq results.

We assessed concordance between snRNA-seq and spatial transcriptomic measurements in two ways. First, we mapped the cell-type assignments from snRNA-seq onto the Visium HD gene expression grids for aHC and pHC, confirming that Visium HD recapitulated the expected spatial organization of major cell classes across hippocampal subfields (Supplementary Fig. S3A). Second, we quantitatively compared gene expression profiles between platforms within matched cell classes. Gene-wise expression levels showed strong concordance between snRNA-seq and Visium HD for excitatory and inhibitory neurons at both hippocampal poles (Exc: aHC: Spearman ρ = 0.42, 95% CI [0.40-0.44]; pHC: ρ = 0.45, 95% CI [0.43-0.46]; Inh: aHC: Spearman ρ = 0.38, 95% CI [0.37-0.40]; pHC: ρ = 0.40, 95% CI [0.38-0.41]; Fig. S3B), with highly significant correlations (p < 2.2 × 10^−16^) and robust agreement under permutation testing. Based on these results, we conclude that Visium HD and snRNA-seq are highly concordant for cross-platform comparisons. Further, Visium HD analysis of the hippocampus confirms the A-P transcriptional gradients in both excitatory and inhibitory neurons that were revealed by snRNA-seq. Next, we sought to link these gene expression gradients to oscillatory fingerprints of successful memory formation by comparing single-cell transcriptional profiles with band-limited power and ripple activity.

### Coupling hippocampal SMEs to gene expression and regulatory programs

For this analysis, we adapted our previously validated within-subject correlation framework^29, 30^ to relate subject-specific SMEs and ripple rates with cell-type-specific pseudobulk gene expression from matching aHC and pHC samples. Analyses were performed separately for excitatory (EXC) and inhibitory (INH) neurons. A two-stage correlation and regression approach identified transcripts whose expression most strongly predicted variance in oscillatory encoding signatures (Supplementary Table S5).

Integrating electrophysiological and transcriptomic data revealed distinct molecular programs associated with encoding-related oscillations in aHC versus pHC. In aHC, delta/theta-associated transcripts were enriched for genes involved in synaptic remodeling (such as *FOSB* and *EGR1*) and chromatin regulation (such as *BCL11A* and *PCGF6*) (Fig. 4A). Genes showing the strongest positive correlations included *CTNNB1*, *RBFOX1*, *DLGAP1*, and *CAMK4* across both excitatory and inhibitory populations, overlapping with delta-related genes previously identified in the temporal pole^29^. In contrast, gamma- and ripple-linked genes in pHC emphasized metabolic, translational and proteostatic programs (e.g., *ATP5PB*, *NDUFB2*, *COX4I1*, *PSMA5*, *USP14*, *EIF4G1*, *EIF4G3*, *PINK1*, *PRKAA2*, and *MTOR,* Supplementary Table S5). Thus, anterior-associated transcripts were enriched for structural and regulatory programs, whereas posterior-associated transcripts were enriched for metabolic and translational programs. Consistent with these trends, posterior enrichment of *KCNQ5*/*KCNIP2*, *SHISA6*/*CACNG8* and *DLGAP1*/*PPFIA2* identified a set of genes linked to resonance, AMPAR-associated signaling, and post-synaptic scaffolding. In contrast, anterior enrichment of *GRM4*, *GRM8*, and *CHRM2* highlighted metabotropic and modulatory signaling pathways. Together, these findings indicate that oscillatory specialization along the hippocampal axis is mirrored by distinct cell-type-specific transcriptional programs.

**Figure 4.**
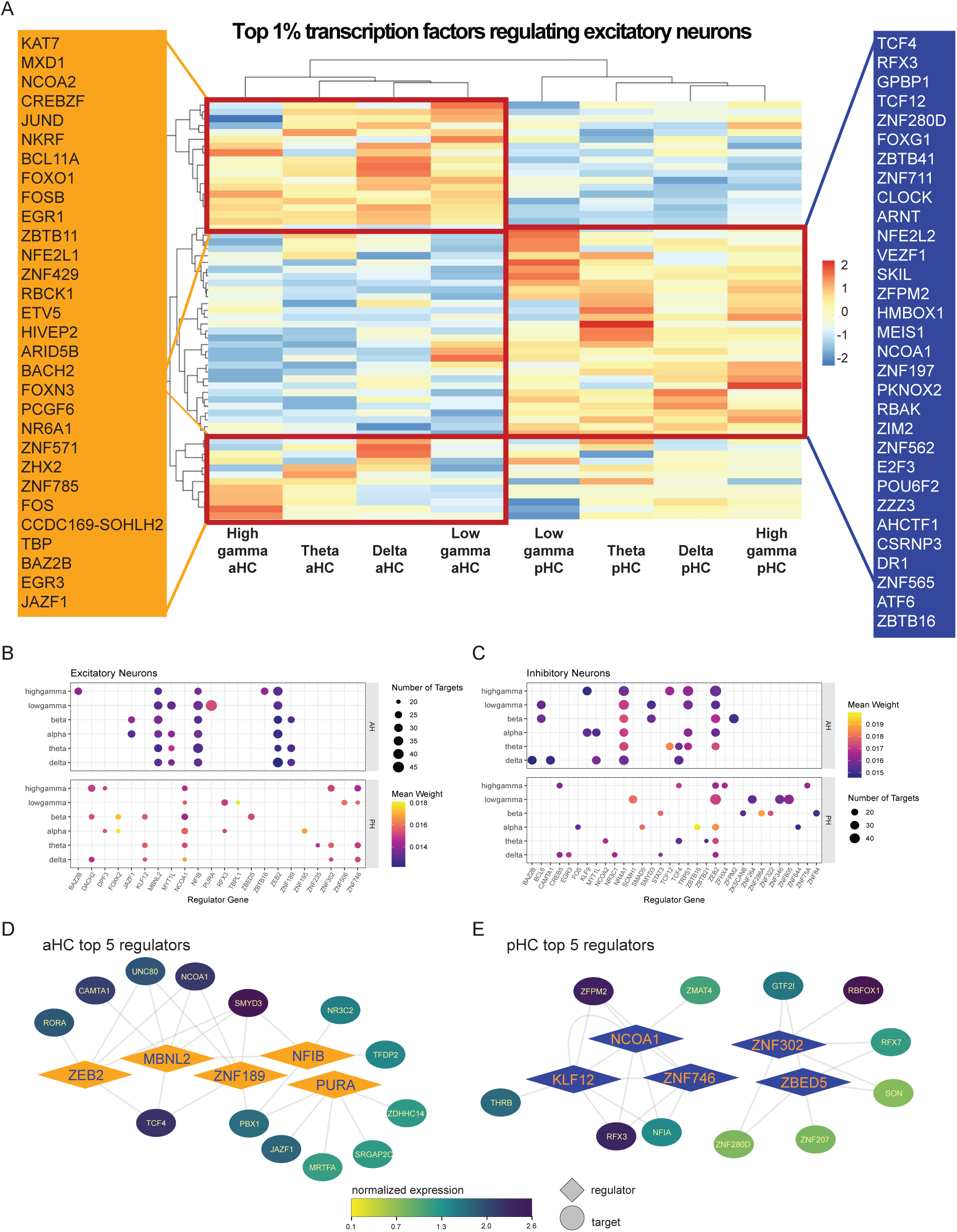
Longitudinal specialization of gene regulatory networks in the human hippocampus. **A-C:** Heatmap (A) and dot plots (B-C) of SME-correlated gene regulatory networks aHC and pHC, shown separately for EXC and INH, GRN hub maps for EXC: aHC (top) vs pHC (bottom) highlighting multiband regulators (*ZEB2*, *NFIB*, *MBNL2*, *ZNF189*, *MYT1L* in aHC; *DACH2*, *ZNF746*, *KLF12*, *NCOA1*, *ZNF302* in pHC). **F,** GRN hub maps for INH: aHC (top) vs pHC (bottom) showing *NR4A1*, *ZEB2*, *TRPS1*, *TCF4*, *KLF9*, *BAZ2B* (aHC) and *ZEB2*, *CREB5*, *STAT3*, *ZNF84/286A/322*, *SCMH1* (pHC). **D-E,** Top 5 dominant excitatory neuron regulatory hubs in aHC (D) and pHC (E). Regulators from GENIE3 using top1% of inferred interactions requiring at least 15 target genes. The top 5 highest expressed genes associated with each hub are displayed. Diamond nodes represent hub regulators, and circular nodes are target genes. Target node color denotes normalized expression across subjects.

### Gene regulatory network (GRN) inference

To determine whether SME-linked transcripts converged into coherent regulatory architectures, we inferred cell-type-specific gene regulatory networks (GRNs) using GENIE3 and identified high-confidence hub regulators for each region and oscillatory band (Fig. 4; Supplementary Table S6).

In excitatory neurons, anterior hippocampal gene-networks were anchored by regulators involved in transcriptional control, chromatin state, and activity-dependent gene expression, including *ZEB2*, *NFIB*, *MBNL2*, *ZNF189*, and *MYT1L*. These hubs were preferentially associated with delta/theta-linked transcripts.

In contrast, posterior excitatory networks were centered on hubs implicated in metabolic regulation and activity-scaling, including *DACH2*, *ZNF746*, *KLF12*, *NCOA1* (SRC-1), and *ZNF302*. Additional posterior hubs (*RFX3*, *ZNF506*, *TBPL1*, *DPF3*) were linked to ciliogenesis and chromatin remodeling (Fig. 4A).

In inhibitory neurons, anterior hippocampal GRNs were similarly enriched for transcriptional and chromatin-associated regulators, including *NR4A1*, *ZEB2*, *TRPS1*, *TCF4*, *KLF9*, and *BAZ2B*. Posterior inhibitory networks, by contrast, were characterized by hubs including *CREB5*, *STAT3*, *ZEB2*, and *SCMH1*, emphasizing energy-responsive transcription and rapid network recruitment.

Across neuronal cell types, SME-linked gene expression and associated GRNs revealed a coherent axis-specific regulatory organization. Anterior networks are enriched for transcriptional and chromatin-centered regulators associated with delta/theta SMEs, whereas posterior networks are enriched for metabolic and activity-scaling regulators associated with gamma- and ripple-related SMEs (Fig. 4A-C). Together with the axis-specific effector gene gradients in excitatory and inhibitory neurons, these findings define a coherent physiological–transcriptional–regulatory organization of neural features supporting human hippocampal encoding.

### Non-neuronal transcriptional and regulatory specialization across the hippocampal axis

Given growing evidence that non-neuronal cells contribute to hippocampal circuit function and memory related plasticity, we assessed whether transcriptional gradients extended beyond neuronal populations. Differential expression analysis across non-neuronal cell classes revealed significant anterior-posterior differences selectively in astrocytes, while other glial populations did not exhibit a regional bias (Supplementary Table S3 and S4). Posterior hippocampal astrocytes showed enrichment of genes associated with synaptic regulation, extracellular matrix organization, and neuromodulatory signaling, including *KCND3*, *EPHA4*, *NCAN*, *CHRDL1* and *AXL* (Supplementary Table S3 and S4). In contrast, astrocytes in the anterior hippocampus did not exhibit a corresponding set of region-biased transcripts.

We next asked whether axis-specific regulatory organization also extended to non-neuronal populations. Oligodendrocyte GRNs exhibited clear anterior-posterior dissociations paralleling those observed in neurons. Anterior oligodendrocyte networks were anchored by transcriptional regulators associated with differentiation, chromatin state, and myelin stabilization including *ZNF652*, *KLF9*, *PBX1*, *ZBTB16*, and *OLIG1*, whereas posterior oligodendrocyte networks were dominated by stress-responsive, immune-metabolic, and mitochondrial regulators including *ST18*, *STAT2*, *IKZF2*, *ZBTB7A*, and *BCL6*. Together, these findings indicate that axis-specific molecular specialization extends beyond neurons to glial populations.

## Discussion

By integrating intrahippocampal iEEG during episodic encoding with hippocampal transcriptomics from the same individuals, we identify a coordinated molecular–physiological organization along the human hippocampal longitudinal axis. Across gene expression, SME–expression coupling, spatial validation, and regulatory network analysis, anterior hippocampus preferentially links mnemonic neurophysiology to modulatory and plasticity-weighted programs, whereas posterior hippocampus links memory processing to high-throughput synaptic and metabolic programs. Our data motivate a framing by which longitudinal regions differ in their preferred operating dynamics: aHC is biased toward neuromodulator-sensitive integration and longer-timescale reconfiguration, whereas pHC is biased toward energetically demanding computations associated with fast oscillations and ripple events.

Prior to incorporation of oscillatory information, we observed pronounced longitudinal transcriptional gradients that were stronger in excitatory than inhibitory neurons. The posterior excitatory program included *GRIA1*, *GABRG2*, *SHISA6*, *CACNG8*, *KCNQ5*, *KCNIP2*, *DLGAP1* and *PPFIA2*, genes linked to glutamatergic signaling, AMPAR-associated function, excitability tuning, and postsynaptic organization - features consistent with rapid circuit dynamics and precise temporal coordination. In contrast, anterior excitatory neurons showed enrichment of *RELN*, *GRM4*, *GRM8*, and *CHRM2*, emphasizing metabotropic, cholinergic modulators and adhesion related programs, consistent with temporal filtering and stable coupling within broader networks^31–34^. Visium HD maps from independent samples recapitulated axis-differing transcripts and their spatial localization, supporting the robustness of these gradients in situ despite expected cross-platform differences in sensitivity and cellular resolution.

Superimposed on these gradients, SME–expression correlation models revealed transcriptional programs that differed across the poles and frequency bands. In aHC, delta/theta SMEs covaried with genes and pathways linked to synaptic remodeling and chromatin/RNA-regulatory function, including *CTNNB1*, *RBFOX1*, *DLGAP1*, *CAMK*4, *BCL11A*, and *PCGF6*, consistent with encoding states that engage activity-dependent plasticity mechanisms^29^. In pHC, gamma- and ripple-related SMEs covaried with metabolic, translational, and proteostatic programs, including genes such as *ATP5PB*, *NDUFB2*, *COX4I1*, *PSMA5*, *USP14*, *EIF4G1*, and *MTOR*, consistent with the energetic and biosynthetic demands of sustaining fast oscillations and ripple-associated bursts and potentially faster synaptic dynamics^20–22, 25^. Together, these relationships suggest that longitudinal specialization is expressed not only in baseline molecular identity but also in how local physiology couples to distinct cellular economies during successful encoding with convergent gene programs across both analyses.

Gene regulatory network inference sharpened these observations by separating “effectors” from candidate “control knobs.” In both excitatory and inhibitory neurons, aHC SME-linked networks were disproportionately organized around hub regulators implicated in transcriptional control, chromatin state, and activity-dependent regulation^35–37^, consistent with slower-timescale tuning of circuit properties. In pHC, networks emphasized hub regulators such as *DACH2*, *KLF12*, *NCOA1*, *ZNF746*, and *ZNF302*, with additional hubs including *RFX3*, *TBPL1*, and *DPF3*, pointing toward metabolic regulation, activity scaling and chromatin-associated adaptation consistent with the energetic demands of high-frequency network activity^38, 39^. This regulator-level dissociation provides a plausible mechanism by which the poles could maintain different operating regimes: effector gradients bias local circuit physiology, while distinct hub architectures set the preferred timescale and mode (e.g., cholinergic tone^34, 40^) through which encoding states are stabilized and adapted across experience.

Long-axis specialization also extended beyond neurons. Posterior astrocytes were enriched for *KCND3*, *EPHA4*, *NCAN*, *CHRDL1*, and *AXL*, implicating astrocyte-associated regulation of excitability, extracellular matrix organization, and synaptic signaling^41–44^. Oligodendrocyte GRNs showed a parallel anterior-posterior dissociation, with anterior networks centered on Z*NF652*, *KLF9*, *PBX1*, *ZBTB16*, and *OLIG1*, emphasizing differentiation and myelin stabilization and posterior networks centered on *ST18*, *STAT2*, *IKZF2*, *ZBTB7A*, and *BCL6*, emphasizing stress-responsive immune–metabolic and mitochondrial regulation. These findings are consistent with growing evidence that glial cells contribute to memory-related plasticity, including oligodendrogenesis-dependent memory consolidation in adult mice^44–47^, and suggest that longitudinal hippocampal specialization involves cell type specific transcriptional organization.

These findings intersect with longstanding accounts of anterior-posterior functional differentiation but extend them by grounding physiological specialization in molecular programs identified from human tissue. We interpret the transcriptional differences across neuronal subtypes and SME-coupled gene programs as converging on a model of a longitudinal gradient in physiological constraints: modulation and plasticity readiness in aHC versus synaptic throughput and energetic capacity in pHC, from which representational differences could emerge as a function of task demands.

At the same time, alternative interpretations remain possible. Rather than reflecting fixed specialization, anterior and posterior hippocampus may participate flexibly in multiple distributed cortical networks, with task demands determining which circuits are engaged. Nevertheless, the convergence across gene expression, SME coupling, spatial validation, and regulatory network analyses provides empirical constraints that can inform and discipline future computational models of longitudinal hippocampal function. More broadly, the axis- and state-specific hub regulators identified here nominate molecular targets that could, in principle, be leveraged for region- and mnemonic-specific neuromodulation strategies

### Limitations and Caveats

Several limitations qualify interpretation. Our cohort comprises patients with refractory epilepsy, and disease processes and antiseizure medications can influence both excitability and transcriptional state. Sample sizes were constrained by clinical tissue availability, but pseudobulk approaches mitigated donor-level variability. Finally, all gene–physiology relationships reported here are correlational; causal tests will require experimental manipulation of oscillatory regimes and/or targeted perturbation of inferred hub regulators, ideally paired with longitudinal molecular readouts. Despite these constraints, the convergence across differential expression, SME–expression coupling, spatial validation, and GRN architecture provides a coherent molecular account of how encoding-related hippocampal physiology is organized along the human long axis.

In addition, the behavioral paradigms used here primarily probe verbal episodic memory, and it remains unclear whether similar molecular-physiological relationships generalize across other hippocampal-dependent domains such as spatial navigation, affective memory, or imagination.

## Methods

### Patient cohort and ethics

Patients with pharmacoresistant temporal lobe epilepsy undergoing presurgical evaluation at the Epilepsy Monitoring Unit at the University of Texas Southwestern Medical Center (Dallas, TX) were recruited for this study. All procedures were approved by the UT Southwestern Institutional Review Board under the IRB protocols STU 092014-026 and 092014-075, and all participants provided written informed consent. Intracranial EEG (iEEG) data were collected from eleven patients who completed at least one episodic memory task during presurgical monitoring. A subset of fourteen patients, including the eleven contributing iEEG data, subsequently underwent surgical resection and contributed hippocampal tissue for (spatial-)transcriptomic analyses (Table S1). One patient did not meet quality criteria for iEEG analysis but was included in transcriptomic analyses, two patients were included for independent spatial transcriptomic validation analyses. Electrode number and placement were determined solely by clinical considerations. Anatomical localization of depth contacts to anterior and posterior hippocampus was verified by expert manual neuroradiological review.

### Memory task design

Participants completed one or both of two verbal episodic memory paradigms during iEEG recording: Associative Recognition (AR) and Free Recall (FR) both previously described^21, 29^, here briefly outlined.

#### Associative Recognition task

In the AR task, participants studied visually presented, semantically unrelated word pairs composed of concrete nouns (3-9 letters), selected from established word association norms. During encoding, 240 word pairs were presented. At retrieval, participants were shown 160 intact pairs, 80 rearranged pairs (previously seen words paired with a different word), and 80 novel pairs. Participants indicated whether each test pair was intact, rearranged, or novel.

Successful encoding was defined as correct identification of intact pairs (“associative hits”), whereas incorrect responses were classified as unsuccessful encoding (“associative misses”). Participants completed a practice block prior to experimental trials, which was excluded from analysis.

#### Free Recall task

In the FR task, participants completed multiple study–test cycles. During each study phase, 12 high-frequency, single-syllable nouns were visually presented sequentially (1.6 s per word), followed by a jittered blank interval (mean 4 s). After presentation of the final word, participants performed a 30 s arithmetic distractor task to reduce rehearsal, followed by a verbal recall period in which they recalled as many items as possible in any order. Each session consisted of 12 study–test cycles and one practice cycle (excluded from analysis). Participants completed between 1 and 9 sessions (median = 2). Behavioral performance was quantified as recall fraction. Only sessions with recall fraction >10% were included.

### iEEG acquisition and preprocessing

Continuous iEEG signals were recorded during task performance. Signals were referenced and preprocessed using established pipelines. Data were epoched from −1.0 s to +1.6 s relative to stimulus onset for ripple analyses and from −0.5 s to +1.6 s for spectral power analyses. Signals were downsampled to 1 kHz for ripple detection and to 250 Hz for spectral analyses.

### Computation of SMEs

SMEs were computed separately for AR and FR tasks. For AR, encoding epochs were categorized based on later associative hits versus misses. For FR, encoding epochs were categorized based on whether items were subsequently recalled or not recalled.

#### Oscillatory power

Band-limited power was computed spanning 2–120 Hz and grouped into six canonical frequency bands: delta (2-4 Hz), theta (4-8 Hz), alpha (8-16 Hz), beta (16-30 Hz), low gamma (30-70 Hz), and high gamma (70-120 Hz).

For each electrode, power was summed across frequency bins within each band, converted to decibel units, and z-scored relative to a prestimulus baseline (−0.5 to 0 s). For each electrode, SMEs were calculated as the difference in mean power between successful and unsuccessful encoding trials. SMEs were then aggregated across electrodes within anterior and posterior hippocampal regions for subject-level analyses.

#### Ripple detection

Putative hippocampal ripples were detected from the 83-117 Hz envelope as previously described^27^. A ripple candidate was defined as a contiguous segment exceeding 2-3 standard deviations above the trial-specific mean for at least 20 ms; events separated by ≤5 ms were merged. Ripple counts were restricted to 0.5–1.6 s poststimulus to avoid stimulus-locked transients. Ripple rate was computed per electrode as the mean number of ripples per trial divided by the analysis window and averaged across electrodes within each hippocampal region. Ripple SMEs were defined as the difference in ripple rate between successful and unsuccessful encoding trials. All oscillatory analyses were implemented in MATLAB using the Lega Lab iEEG toolbox.

### Hippocampal tissue collection

Hippocampal tissue was obtained from patients undergoing anterior temporal lobectomy as part of clinical care. After resection of lateral temporal cortex, the hippocampus was removed using an en bloc surgical technique preserving its longitudinal axis. Only specimens measuring at least 3.5 cm along the anterior-posterior axis were included. Approximately 0.5 cm segments were sampled from the anterior and posterior poles; intervening tissue was submitted for clinical pathology. Anterior samples were defined as tissue anterior to the uncal notch. Tissue was immediately placed in ice-cold Neurobasal-A medium, transported to the laboratory within 20 minutes, washed in PBS, and flash-frozen in liquid nitrogen.

### snRNA-seq

snRNA-seq was performed using the ScaleBio (now 10X Genomics) platform^48^. Nuclei isolation followed an adapted protocol based on established methods for human hippocampal tissue and 10x Genomics demonstrated protocols (CG000375). Briefly, frozen tissue was homogenized using a Dounce homogenizer in lysis buffer, incubated on ice, and centrifuged at 500 x g at 4°C. Lysis was repeated for a total of 10 minutes. Pellets were washed in nuclei suspension buffer, filtered through a 40 μm Flowmi filter, and purified using sucrose gradient centrifugation (13,000 x g, 45 min, 4°C). Nuclei were washed, and fixed following ScaleBio nuclei fixation kit. Nuclei were manually counted using Hoechst nuclear stain. Samples were adjusted to 5000 nuclei/μL and processed according to the ScaleBio protocol. Pooled cDNA was amplified, indexed, and purified using SPRIselect beads. Libraries were sequenced on an Illumina NovaSeq 6000, with one lane per pooled sample. snRNA-seq preprocessing and annotationRaw sequencing data were obtained as BCL files from the McDermott sequencing core at UT Southewestern. BCL files were demultiplexed using STARsolo. Samples were integrated using Seurat’s integration approach and integrated datasets were further clustered and visualized (RunUMAP) using Uniform manifold approximation and projection embeddings (UMAP). Cluster-specific gene markers were identified (FindMarkers) and significant marker genes were enriched using Fisher exact test for cell-type markers defined in Brain Initiative Cell Census and our previously published human hippocampal reference atlas^28^. Cell types were also confirmed by expression of canonical marker genes.

#### Cell-type composition analysis

Cluster-specific nucleus counts were modeled using Poisson generalized linear mixed-effects models with hippocampal region as a fixed effect, donor as a random intercept, and an offset for the total number of nuclei per donor and region. This approach tested for regional differences in relative cell-type abundance while accounting for inter-individual variability (Fig. S2, Table S4).

### Visium HD spatial transcriptomics

#### RNA quality assessment

Formalin-fixed, paraffin-embedded (FFPE) hippocampal sections previously stained with H&E were assessed for RNA quality following 10X Genomics guidelines (CG000684). Coverslips were removed using xylene, tissue was scraped, and RNA was extracted using the RNeasy FFPE Kit (Qiagen). RNA integrity was assessed using the DV200 metric on an Agilent TapeStation; all samples had DV200 >30%.

#### Imaging and Visium HD workflow

Sequential tissue sections were imaged using a Nikon AXR confocal microscope equipped with an Fi3 color camera. Whole-slide brightfield images were acquired at 20x magnification. Visium HD libraries were prepared following 10X Genomics guidelines (CG000685) using the Human Transcriptome Probe Kit v2. Probe hybridization was performed overnight, followed by ligation and CytAssist-enabled probe transfer. cDNA was amplified, indexed, and purified using SPRIselect beads. Library quality was assessed using a High Sensitivity DNA assay on an Agilent TapeStation. Libraries were sequenced on an Illumina NovaSeq 6000, with one lane per sample.

Differential Gene Expression tests were performed using DESeq2 based pseudobulk approach across hippocampal axis per cell type. Significantly differentially expressed genes (DEGs) were identified using absolute log2fold change of ≥ 0.333 (26% change in expression) and a false discovery rate (FDR) ≤ 0.05.

Gene regulatory networks (GRNs) were inferred separately for anterior and posterior hippocampal excitatory and inhibitory neurons using the GENIE3 algorithm^49, 50^, which employs ensemble tree-based regression to predict regulator–target relationships from gene expression data. Regulatory interactions were ranked by GENIE3 importance scores, and high-confidence edges were defined as those within the top 1% of interaction weights. Hub regulators were identified as transcription factors associated with at least 15 inferred target genes. For visualization, the five highest-ranked hub regulators from anterior and posterior excitatory neuron networks were selected, and the five most highly expressed target genes associated with each regulator were retained. Network visualization was performed in Cytoscape^51^, with regulator nodes displayed separately from target genes and target node color scaled according to average normalized gene expression across subjects.

### Experimental design and statistical considerations

Sample sizes were determined by tissue availability and clinical constraints. No statistical methods were used to predetermine sample size. Samples were not randomized. Sex chromosomes were excluded from gene expression analyses, and sex was included as a covariate where appropriate.

## Acknowledgements

We are deeply grateful to the patients and their families who participated in these research studies, without whose selfless generosity this work would not have existed. We thank the O’Donnell Brain Institute at UTSW and the members of UT Southwestern Neuroscience and Neurosurgery departments for their assistance and support with this research.

## Funding

National Institutes of Health grant NS126143 (BCL, GK)

## Author contributions

Conceptualization: MD, BCL, GK

Methodology: MD, TT, BCL, GK

Investigation: MD, HM, TT, AF

Formal analysis: MD, TT, AK

Resources: AK, BCL, GK

Visualization: MD, TT, AK

Supervision: BCL, GK

Writing – original draft: MD

Writing – review & editing: MD, HM, AF, BCL, GK

## Competing interests

The authors declare that they have no competing interests.

---

**Supplementary Figure S1.**
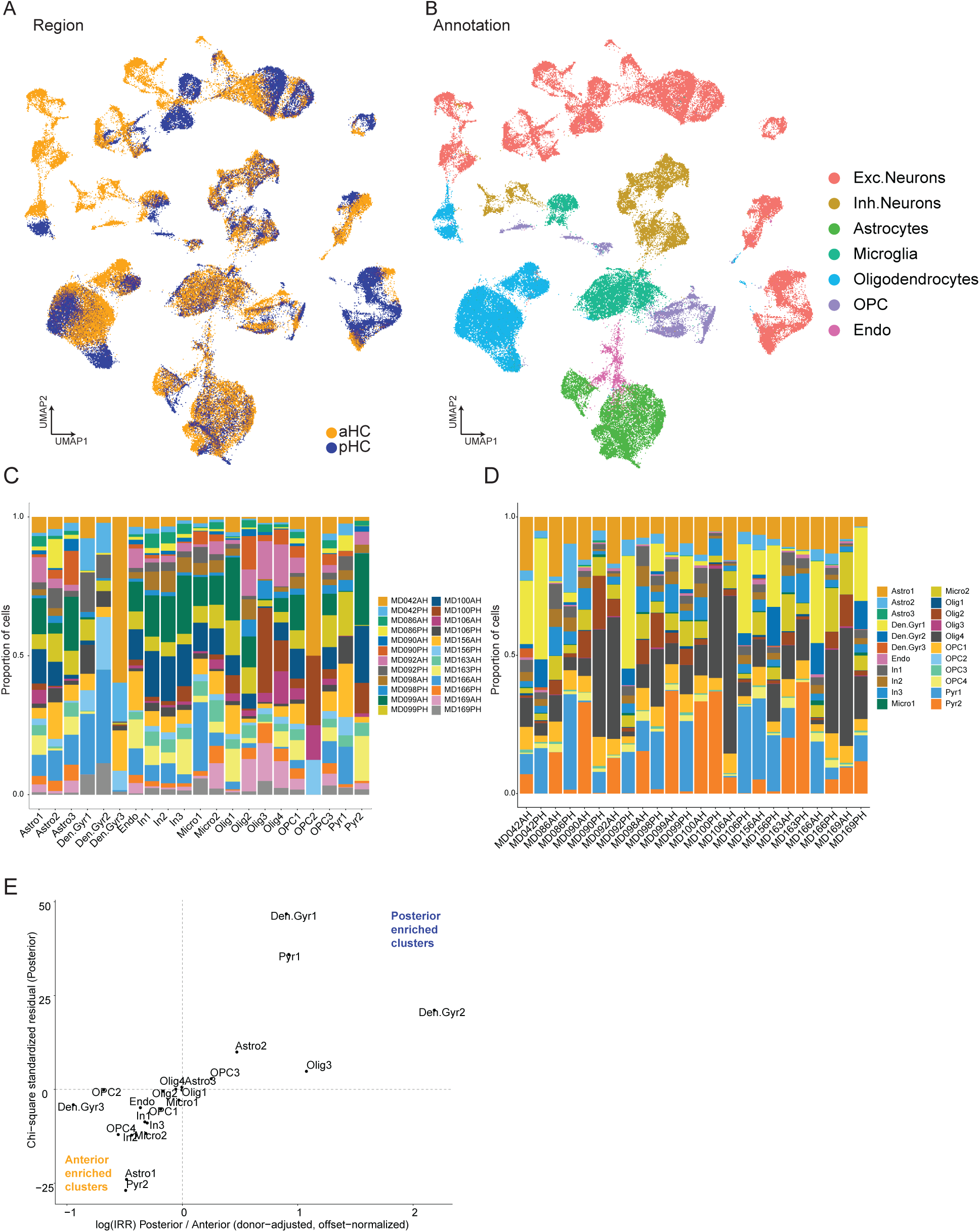
Cell-type composition and sample contribution across anterior and posterior hippocampus. **A:** UMAP embedding of all nuclei colored by hippocampal pole (anterior, aHC; posterior, pHC), showing intermixing of nuclei across regions and absence of large-scale segregation driven by sampling location. **B:** UMAP embedding of the same nuclei colored by major cell classes, including excitatory and inhibitory neurons, oligodendrocytes, oligodendrocyte precursor cells, astrocytes, microglia, and endothelial cells, annotated using canonical marker genes. **C:** Bar plot showing the proportional contribution of samples to each cell-type cluster (columns represent clusters), demonstrating balanced representation of anterior and posterior hippocampal samples across clusters. **D:** Bar plot showing the distribution of cell-type clusters within each individual sample (columns represent samples), indicating consistent cellular composition across donors and hippocampal regions. **E:** Scatter plot comparing χ² residuals from contingency analyses (posterior column) with donor-adjusted log incidence rate ratios (IRRs) derived from mixed-effects models normalizing for total nuclei per donor and region. Directional concordance was observed for 20 of 22 clusters; the two discordant clusters showed negligible effect sizes (logIRR ≈ 0 and χ² residual ≈ 0), indicating minimal biological relevance and supporting the absence of systematic compositional bias between anterior and posterior hippocampus.

**Supplementary Figure S2.**
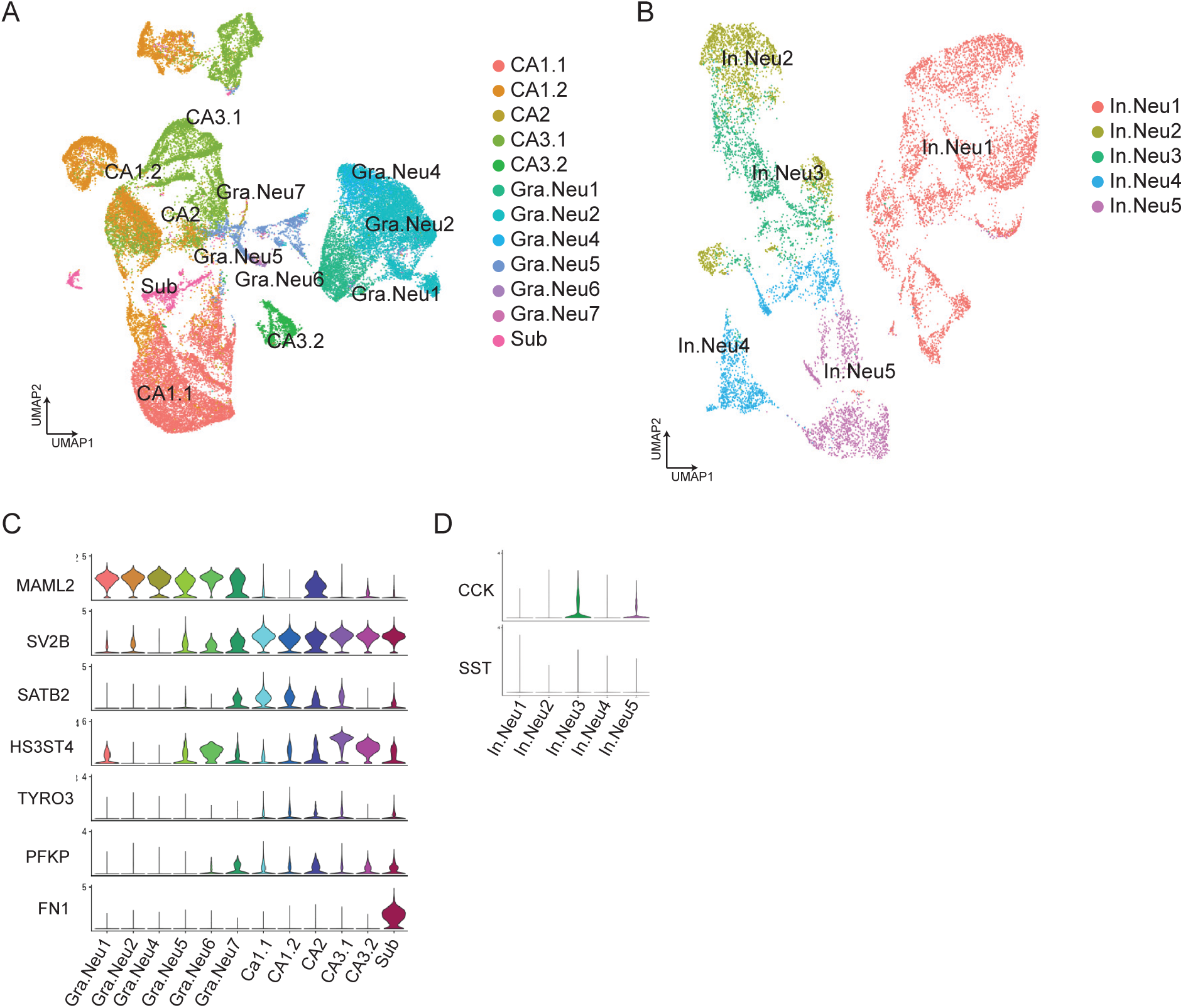
Subclustering of excitatory and inhibitory neurons and marker-gene validation. **A**: UMAP embedding of excitatory neuronal nuclei following subclustering, showing resolution of excitatory subclasses. **B:** UMAP embedding of inhibitory neuronal nuclei following subclustering, showing resolution of inhibitory subclasses. **C:** Violin plots showing expression of subclass marker genes across neuronal subclusters (MAML2, SV2B, SATB2, HS3ST4, TYRO3, PFKP, FN1). **D**: Violin plots showing expression of interneuron marker genes *SST* and *CCK* across inhibitory subclusters, supporting annotation of major interneuron subclasses.

**Supplementary Figure S3.**
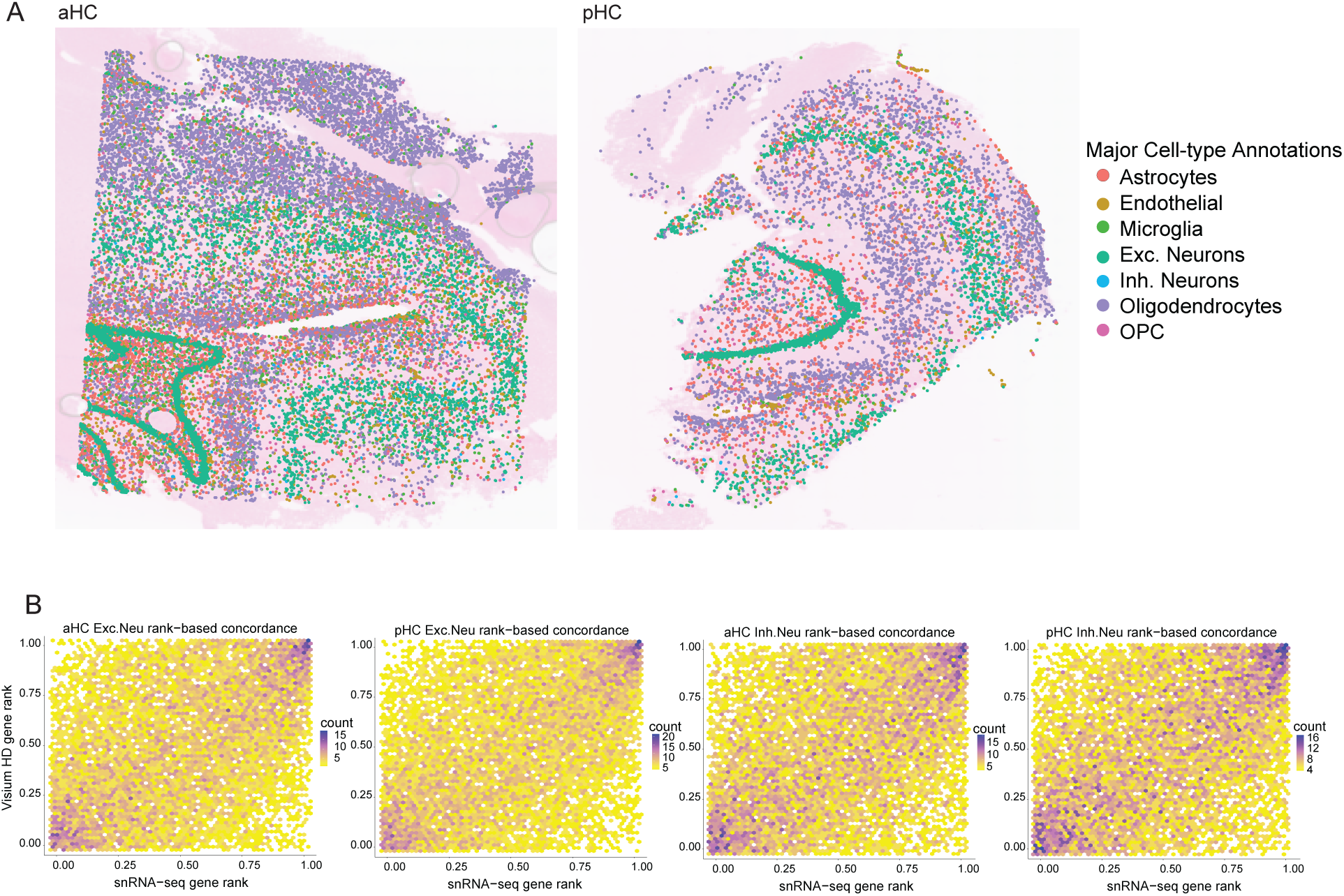
Spatial transcriptomic mapping and cross-platform concordance. **A**: Spatial DimPlot showing the distribution of major cell classes mapped onto Visium HD sections from anterior hippocampus (aHC; left) and posterior hippocampus (pHC; right). Cell-type assignments were transferred from the snRNA-seq reference and recapitulate expected laminar and regional organization across hippocampal subfields. **B**: Rank-based concordance between Visium HD and snRNA-seq gene expression estimates. Scatter plots show gene-wise correspondence between platforms within matched hippocampal poles for excitatory (first two) and inhibitory (second two) neuronal populations, shown separately for anterior (aHC) and posterior (pHC) hippocampus. (Exc: aHC: Spearman ρ = 0.42, 95% CI [0.40-0.44]; pHC: ρ = 0.45, 95% CI [0.43-0.46]; Inh: aHC: Spearman ρ = 0.38, 95% CI [0.37-0.40]; pHC: ρ = 0.40, 95% CI [0.38-0.41]; p < 2.2 × 10^−16^ for all comparisons).

## Notes

### Competing Interest Statement

The authors have declared no competing interest.

## REFERENCES

1. Kapardi M, Kavitha A, editors. Functional connectivity assessment for episodic memory. Proceedings of 2017 IEEE 16th International Conference on Cognitive Informatics and Cognitive Computing, ICCI*CC 2017; 2017.

2. Bettcher BM. Memory, Episodic. Encyclopedia of the Neurological Sciences2014. p. 1039–41.

3. Ranganath C. Episodic memory. The Oxford Handbook of Human Memory: Foundations and Applications2024. p. 151–71.

4. Li JS, Chao YS. Electrolytic lesions of dorsal CA3 impair episodic-like memory in rats. Neurobiology of Learning and Memory. 2008;89(2):192–8. doi: 10.1016/j.nlm.2007.06.006.

5. Raud L, Sneve MH, Vidal-Piñeiro D, Sørensen Ø, Folvik L, Ness HT, Mowinckel AM, Grydeland H, Walhovd KB, Fjell AM. Hippocampal-cortical functional connectivity during memory encoding and retrieval. NeuroImage. 2023;279. doi: 10.1016/j.neuroimage.2023.120309.

6. Lega BC, Jacobs J, Kahana M. Human hippocampal theta oscillations and the formation of episodic memories. Hippocampus. 2012;22(4):748–61. Epub 20110427. doi: 10.1002/hipo.20937. PubMed PMID: 21538660.

7. Lega B, Burke J, Jacobs J, Kahana MJ. Slow-Theta-to-Gamma Phase-Amplitude Coupling in Human Hippocampus Supports the Formation of New Episodic Memories. Cerebral cortex. 2016;26(1):268–78. Epub 20141014. doi: 10.1093/cercor/bhu232. PubMed PMID: 25316340; PMCID: PMC4677977.

8. Herweg NA, Solomon EA, Kahana MJ. Theta Oscillations in Human Memory. Trends Cogn Sci. 2020;24(3):208–27. Epub 20200203. doi: 10.1016/j.tics.2019.12.006. PubMed PMID: 32029359; PMCID: PMC8310425.

9. Sederberg PB, Schulze-Bonhage A, Madsen JR, Bromfield EB, McCarthy DC, Brandt A, Tully MS, Kahana MJ. Hippocampal and neocortical gamma oscillations predict memory formation in humans. Cerebral cortex. 2007;17(5):1190–6. Epub 20060710. doi: 10.1093/cercor/bhl030. PubMed PMID: 16831858.

10. Burke JF, Sharan AD, Sperling MR, Ramayya AG, Evans JJ, Healey MK, Beck EN, Davis KA, Lucas TH, 2nd, Kahana MJ. Theta and high-frequency activity mark spontaneous recall of episodic memories. The Journal of neuroscience : the official journal of the Society for Neuroscience. 2014;34(34):11355–65. doi: 10.1523/JNEUROSCI.2654-13.2014. PubMed PMID: 25143616; PMCID: PMC4138344.

11. Norman Y, Yeagle EM, Khuvis S, Harel M, Mehta AD, Malach R. Hippocampal sharp-wave ripples linked to visual episodic recollection in humans. Science. 2019;365(6454). doi: 10.1126/science.aax1030. PubMed PMID: 31416934.

12. Duan QT, Dai L, Wang LK, Shi XJ, Chen X, Liao X, Zhang CQ, Yang H. Hippocampal ripples correlate with memory performance in humans. Brain Res. 2023;1810:148370. Epub 20230418. doi: 10.1016/j.brainres.2023.148370. PubMed PMID: 37080267.

13. Freelin A, Wolfe C, Lega B. Models of human hippocampal specialization: a look at the electrophysiological evidence. Trends in Cognitive Sciences. 2024. doi: 10.1016/j.tics.2024.11.009.

14. Moser MB, Moser EI. Functional differentiation in the hippocampus. Hippocampus. 1998;8(6):608–19. doi: 10.1002/(SICI)1098-1063(1998)8:6<608::AID-HIPO3>3.0.CO;2-7; PMCID: 9882018.

15. Small SA, Nava AS, Perera GM, DeLaPaz R, Mayeux R, Stern Y. Circuit mechanisms underlying memory encoding and retrieval in the long axis of the hippocampal formation. Nature neuroscience. 2001;4(4):442–9. doi: 10.1038/86115. PubMed PMID: 11276237.

16. Persson J, Söderlund H. Hippocampal hemispheric and long-axis differentiation of stimulus content during episodic memory encoding and retrieval: An activation likelihood estimation meta-analysis. Hippocampus. 2015;25(12):1614–31. doi: 10.1002/hipo.22482.

17. Poppenk J, Evensmoen HR, Moscovitch M, Nadel L. Long-axis specialization of the human hippocampus. Trends in Cognitive Sciences. 2013;17(5):230–40. doi: 10.1016/j.tics.2013.03.005.

18. Strange BA, Witter MP, Lein ES, Moser EI. Functional organization of the hippocampal longitudinal axis. Nature Reviews Neuroscience. 2014;15(10):655–69. doi: 10.1038/nrn3785.

19. Fanselow MS, Dong HW. Are the dorsal and ventral hippocampus functionally distinct structures? Neuron. 2010;65(1):7–19. Epub 2010/02/16. doi: S0896-6273(09)00947-7 [pii] 10.1016/j.neuron.2009.11.031. PubMed PMID: 20152109; PMCID: 2822727.

20. Goyal A, Miller J, Qasim SE, Watrous AJ, Zhang H, Stein JM, Inman CS, Gross RE, Willie JT, Lega B, Lin JJ, Sharan A, Wu C, Sperling MR, Sheth SA, McKhann GM, Smith EH, Schevon C, Jacobs J. Functionally distinct high and low theta oscillations in the human hippocampus. Nature Communications. 2020;11(1). doi: 10.1038/s41467-020-15670-6.

21. Kota S, Rugg MD, Lega BC. Hippocampal Theta Oscillations Support Successful Associative Memory Formation. The Journal of neuroscience : the official journal of the Society for Neuroscience. 2020;40(49):9507–18. Epub 20201106. doi: 10.1523/jneurosci.0767-20.2020. PubMed PMID: 33158958; PMCID: PMC7724134.

22. Lin JJ, Rugg MD, Das S, Stein J, Rizzuto DS, Kahana MJ, Lega BC. Theta band power increases in the posterior hippocampus predict successful episodic memory encoding in humans. Hippocampus. 2017;27(10):1040–53. Epub 20170630. doi: 10.1002/hipo.22751. PubMed PMID: 28608960; PMCID: PMC6517838.

23. Lin JJ, Umbach G, Rugg MD, Lega B. Gamma oscillations during episodic memory processing provide evidence for functional specialization in the longitudinal axis of the human hippocampus. Hippocampus. 2019;29(2):68–72. Epub 20181105. doi: 10.1002/hipo.23016. PubMed PMID: 30394594; PMCID: PMC6519081.

24. Wang DX, Schmitt K, Seger S, Davila CE, Lega BC. Cross-regional phase amplitude coupling supports the encoding of episodic memories. Hippocampus. 2021;31(5):481–92. doi: 10.1002/hipo.23309.

25. To TV, Wang DX, Wolfe CB, Lega BC. Neurophysiological evidence of human hippocampal longitudinal differentiation in associative memory. Nat Commun. 2025;16(1):6845. Epub 20250725. doi: 10.1038/s41467-025-61464-z. PubMed PMID: 40715077; PMCID: PMC12297184.

26. Zhang H, Skelin I, Ma S, Paff M, Mnatsakanyan L, Yassa MA, Knight RT, Lin JJ. Awake ripples enhance emotional memory encoding in the human brain. Nature Communications. 2024;15(1):215. doi: 10.1038/s41467-023-44295-8.

27. Seger S, Ergit E, Arya S, Lega B. Precise temporal dynamics of ripple events support order memory in human hippocampal-cortical circuits. Proc Natl Acad Sci U S A. 2025;122(44):e2422266122. Epub 20251027. doi: 10.1073/pnas.2422266122. PubMed PMID: 41144661; PMCID: PMC12595444.

28. Ayhan F, Kulkarni A, Berto S, Sivaprakasam K, Douglas C, Lega BC, Konopka G. Resolving cellular and molecular diversity along the hippocampal anterior-to-posterior axis in humans. Neuron. 2021;109(13):2091–105.e6. Epub 2021/05/30. doi: 10.1016/j.neuron.2021.05.003. PubMed PMID: 34051145; PMCID: PMC8273123.

29. Berto S, Fontenot MR, Seger S, Ayhan F, Caglayan E, Kulkarni A, Douglas C, Tamminga CA, Lega BC, Konopka G. Gene-expression correlates of the oscillatory signatures supporting human episodic memory encoding. Nature neuroscience. 2021;24(4):554–64. Epub 2021/03/10. doi: 10.1038/s41593-021-00803-x. PubMed PMID: 33686299; PMCID: PMC8016736.

30. Berto S, Wang GZ, Germi J, Lega BC, Konopka G. Human Genomic Signatures of Brain Oscillations During Memory Encoding. Cerebral cortex. 2018;28(5):1733–48. Epub 2017/04/07. doi: 10.1093/cercor/bhx083. PubMed PMID: 28383644; PMCID: PMC5907355.

31. Pujadas L, Gruart A, Bosch C, Delgado L, Teixeira CM, Rossi D, de Lecea L, Martínez A, Delgado-García JM, Soriano E. Reelin regulates postnatal neurogenesis and enhances spine hypertrophy and long-term potentiation. The Journal of neuroscience : the official journal of the Society for Neuroscience. 2010;30(13):4636–49. doi: 10.1523/jneurosci.5284-09.2010. PubMed PMID: 20357114; PMCID: PMC6632327.

32. Kitamura T, Pignatelli M, Suh J, Kohara K, Yoshiki A, Abe K, Tonegawa S. Island cells control temporal association memory. Science. 2014;343(6173):896–901. Epub 20140123. doi: 10.1126/science.1244634. PubMed PMID: 24457215; PMCID: PMC5572219.

33. Kryszkowski W, Boczek T. The G Protein-Coupled Glutamate Receptors as Novel Molecular Targets in Schizophrenia Treatment-A Narrative Review. J Clin Med. 2021;10(7). Epub 20210402. doi: 10.3390/jcm10071475. PubMed PMID: 33918323; PMCID: PMC8038150.

34. Teles-Grilo Ruivo LM, Mellor JR. Cholinergic modulation of hippocampal network function. Front Synaptic Neurosci. 2013;5:2. Epub 20130730. doi: 10.3389/fnsyn.2013.00002. PubMed PMID: 23908628; PMCID: PMC3726829.

35. Yap EL, Greenberg ME. Activity-Regulated Transcription: Bridging the Gap between Neural Activity and Behavior. Neuron. 2018;100(2):330–48. doi: 10.1016/j.neuron.2018.10.013. PubMed PMID: 30359600; PMCID: PMC6223657.

36. Tyssowski KM, Gray JM. The neuronal stimulation-transcription coupling map. Current opinion in neurobiology. 2019;59:87–94. Epub 20190601. doi: 10.1016/j.conb.2019.05.001. PubMed PMID: 31163285; PMCID: PMC6885097.

37. Flavell SW, Greenberg ME. Signaling mechanisms linking neuronal activity to gene expression and plasticity of the nervous system. Annu Rev Neurosci. 2008;31:563–90. doi: 10.1146/annurev.neuro.31.060407.125631. PubMed PMID: 18558867; PMCID: PMC2728073.

38. Kann O, Huchzermeyer C, Kovacs R, Wirtz S, Schuelke M. Gamma oscillations in the hippocampus require high complex I gene expression and strong functional performance of mitochondria. Brain. 2011;134(Pt 2):345–58. Epub 20101222. doi: 10.1093/brain/awq333. PubMed PMID: 21183487.

39. Kann O. The energy demand of fast neuronal network oscillations: insights from brain slice preparations. Front Pharmacol. 2011;2:90. Epub 20120110. doi: 10.3389/fphar.2011.00090. PubMed PMID: 22291647; PMCID: PMC3254178.

40. Hasselmo ME, Sarter M. Modes and models of forebrain cholinergic neuromodulation of cognition. Neuropsychopharmacology. 2011;36(1):52–73. Epub 20100728. doi: 10.1038/npp.2010.104. PubMed PMID: 20668433; PMCID: PMC2992803.

41. Santello M, Toni N, Volterra A. Astrocyte function from information processing to cognition and cognitive impairment. Nature neuroscience. 2019;22(2):154–66. Epub 20190121. doi: 10.1038/s41593-018-0325-8. PubMed PMID: 30664773.

42. Allen NJ, Lyons DA. Glia as architects of central nervous system formation and function. Science. 2018;362(6411):181–5. doi: 10.1126/science.aat0473. PubMed PMID: 30309945; PMCID: PMC6292669.

43. Araque A, Perea G. Glial modulation of synaptic transmission in culture. Glia. 2004;47(3):241–8. PubMed PMID: 6784.

44. Gibson EM, Purger D, Mount CW, Goldstein AK, Lin GL, Wood LS, Inema I, Miller SE, Bieri G, Zuchero JB, Barres BA, Woo PJ, Vogel H, Monje M. Neuronal activity promotes oligodendrogenesis and adaptive myelination in the mammalian brain. Science. 2014;344(6183):1252304. Epub 20140410. doi: 10.1126/science.1252304. PubMed PMID: 24727982; PMCID: PMC4096908.

45. Steadman PE, Xia F, Ahmed M, Mocle AJ, Penning ARA, Geraghty AC, Steenland HW, Monje M, Josselyn SA, Frankland PW. Disruption of Oligodendrogenesis Impairs Memory Consolidation in Adult Mice. Neuron. 2020;105(1):150–64.e6. Epub 20191118. doi: 10.1016/j.neuron.2019.10.013. PubMed PMID: 31753579; PMCID: PMC7579726.

46. Monje M. Myelin Plasticity and Nervous System Function. Annu Rev Neurosci. 2018;41:61–76. doi: 10.1146/annurev-neuro-080317-061853. PubMed PMID: 29986163.

47. Fields RD. A new mechanism of nervous system plasticity: activity-dependent myelination. Nature Reviews Neuroscience. 2015;16(12):756–67. doi: 10.1038/nrn4023.

48. O’Huallachain M, Bava F-A, Shen M, Dallett C, Paladugu S, Samusik N, Yu S, Hussein R, Hillman GR, Higgins S, Lou M, Trejo A, Qin L, Tai YC, Kinoshita SM, Jager A, Lashkari D, Goltsev Y, Ozturk S, Nolan GP. Ultra-high throughput single-cell analysis of proteins and RNAs by split-pool synthesis. Communications Biology. 2020;3(1):213. doi: 10.1038/s42003-020-0896-2.

49. Huynh-Thu VA, Irrthum A, Wehenkel L, Geurts P. Inferring Regulatory Networks from Expression Data Using Tree-Based Methods. PloS one. 2010;5(9):e12776. doi: 10.1371/journal.pone.0012776.

50. Huynh-Thu VA, Geurts P. dynGENIE3: dynamical GENIE3 for the inference of gene networks from time series expression data. Sci Rep. 2018;8(1):3384. Epub 20180221. doi: 10.1038/s41598-018-21715-0. PubMed PMID: 29467401; PMCID: PMC5821733.

51. Shannon P, Markiel A, Ozier O, Baliga NS, Wang JT, Ramage D, Amin N, Schwikowski B, Ideker T. Cytoscape: a software environment for integrated models of biomolecular interaction networks. Genome Res. 2003;13(11):2498–504. doi: 10.1101/gr.1239303. PubMed PMID: 14597658; PMCID: PMC403769.

